# Vacuolar invertase activity shapes photosynthetic stress response of *Arabidopsis thaliana* and stabilizes central energy supply

**DOI:** 10.1101/168617

**Authors:** Jakob Weiszmann, Lisa Fürtauer, Wolfram Weckwerth, Thomas Nägele

## Abstract

Stabilization of the central carbohydrate and energy metabolism plays a key role in plant stress response. As the primary photosynthetic products, carbohydrates are substrate for numerous metabolic and stress-protective reactions. Further, they have been shown to be involved in diverse signalling processes finally affecting and regulating plant stress response on a whole plant level. Sucrose metabolism is known to be central to many stress-related processes and enzymes catalysing its biosynthesis, transport and degradation have been shown to significantly impact stress resistance and acclimation output. However, due to the cyclic structure of sucrose metabolism involving sucrose cleavage in multiple compartments as well as energy-demanding re-synthesis via hexose phosphorylation, it is challenging to derive an unambiguous picture of its contribution to stress reactions. In the present study, a combined stress experiment comprising cold and high-light identified metabolism of sucrose and fumaric acid to significantly separate the stress response of a cold susceptible and a tolerant natural accession of *Arabidopsis thaliana*. Kinetic modelling and simulation of subcellular rates of invertase-driven sucrose cleavage revealed a contrasting picture between the susceptible and the tolerant accession pointing to an important role of vacuolar invertase during initial stress response. Using a T-DNA insertion mutant with a dramatically reduced invertase activity provided evidence for a central role of the enzyme in stabilizing photosynthesis and the central energy metabolism during freezing and high-light stress. Reducing vacuolar invertase activity to about 3% of the wild type resulted in a strong increase of ADP and ATP levels indicating a severe effect on cytosolic and plastidial energy balance. Together with a significant decrease of maximum quantum yield of photosystem II (Fv/Fm) these results suggest that vacuolar invertase activity stabilizes cytosolic energy metabolism by supplying hexose equivalents being phosphorylated in the cytosol. Finally, the accompanying ATP consumption is essential for cytosolic phosphate balance which directly affects photosynthetic performance by the supply of ADP being crucial for photosynthetic ATP production.

## Introduction

Numerous abiotic stress conditions severely affect plant growth and reproductive success. Water deficit and drought, cold and light stress affect a high proportion of global land area and, hence, also plant growth and productivity (Cramer et al., 2011). The abiotic stress response of higher plants is characterized by a comprehensive reprogramming of molecular, biochemical and physiological processes enabling plants to survive and acclimate to a changing environment. While our understanding of plant stress response and acclimation has tremendously improved, e.g. in context of transcriptional regulation (Park et al., 2015), signalling processes (Baxter et al., 2014) or physiological response (Theocharis et al., 2012), the predictive power of current models on abiotic stress response is strongly limited. Amongst others, this limitation is due to a highly compartmentalized and interlaced regulatory network comprising a multitude of regulatory feedback loops and fast-reacting signalling compounds, e.g. reactive oxygen species (ROS), affecting numerous molecular processes (Carmody et al., 2016).

Complexity further increases if response to a combination of (abiotic) stress factors is analysed as they occur in the field. Due to synergistic effects and the high plasticity of plant stress response this frequently leads to observations being difficult to interpret. However, particularly with regard to natural variation of stress response or climate change effects it stands to reason to consider such stress combinations (Ahuja et al., 2010). Previous studies have proven usefulness of combined stress studies. For example, the analysis of transcriptome changes in 10 ecotypes of *Arabidopsis thaliana* under cold, heat, high-light, salt and flagellin treatments as well as under combinations of those revealed that more than 60% of transcriptome changes in response to double stress treatments were not predictable from single stress experiments (Rasmussen et al., 2013). Further, as outlined previously, multiple stresses can interact negatively or positively with regard to plant performance (Suzuki et al., 2014). For example, under combined drought and heat stress growth of *Arabidopsis* was significantly reduced, and additive effects on plant performance could be separated from stress-specific effects (Vile et al., 2012). In contrast, the combination of ozone and drought stress dramatically cancelled effects which were observed under ozone and drought stress alone (Iyer et al., 2013). Dampening of ozone response was discussed to occur due to drought-induced stomatal closure.

Also the combination of low temperature and high-light intensities represents a common field condition (Demmig-Adams et al., 2012). Low temperature has been shown to significantly affect performance and distribution of plants (Boyer, 1982; Stitt and Hurry, 2002). Following exposure to low but non-freezing temperature, many herbaceous plant species, including *Arabidopsis thaliana*, can increase their freezing tolerance in a complex process termed cold acclimation. For a collection of more than 70 natural accessions of *Arabidopsis* from across its native Western Eurasian range a significantly positive correlation between freezing tolerance and latitude of origin has been found (Zhen and Ungerer, 2008), which makes it an attractive system to study cold stress reaction and acclimation. Many studies have made use of the natural genetic variation of freezing tolerance and cold acclimation/deacclimation in *Arabidopsis* providing evidence for a crucial role of the central carbohydrate metabolism within these processes (see e.g. (Hannah et al., 2006; Davey et al., 2009; Nagler et al., 2015; Zuther et al., 2015; Juszczak et al., 2016). A tightly regulated interaction of carbohydrate metabolism with the TCA cycle, amino acid metabolism and flavonoid biosynthesis under cold and high-light conditions has been observed previously (Doerfler et al., 2013). Additionally, subcellular distribution and transport of carbohydrates between plastids, cytosol and vacuole has been shown to be crucially involved in stress-induced reprogramming and stabilisation of metabolism (Wormit et al., 2006; Nägele and Heyer, 2013; Fürtauer et al., 2016; Hoermiller et al., 2017). This immediately implies a strong correlation between cold stress response, photosynthetic performance and light regimes. Previous studies have shown that light is required for enhanced freezing tolerance (Levitt, 1980) indicating a crucial role during cold acclimation (Wanner and Junttila, 1999). Comparing a combined cold/light with a cold/dark treatment revealed that expression of cold-responsive genes was only enhanced in combination with light (Soitamo et al., 2008). While these observations show the dependence of cold stress and acclimation response on a minimum of light, excess light at low temperature leads to an imbalance of absorbed light energy by photochemical reactions and energy utilisation in metabolism. This results in photoinhibition and ROS formation in the thylakoids. Photoinhibition results from imbalance between photodamage of PSII and its repair. As recently summarized and discussed, low temperature stress inhibits the repair of PSII while it does not affect its photodamage (Szymańska et al., 2017). In contrast, high-light intensities cause a net loss of PSII activity although repair of PSII *in vivo* is very efficient over a large range of light intensities (Roach and Krieger-Liszkay, 2014). Consequently, merging high-light with low temperature stress results in a combination of PSII damage and reduced repair activity.

In the present study we exposed two natural accessions with differential cold tolerance and of different geographical origin to (i) cold stress and (ii) combined cold/high-light stress. The analysis of subcellular primary metabolism revealed an interplay of fumaric acid metabolism as well as cytosolic and vacuolar invertase activity separating the metabolic stress response of both accessions. Finally, the contribution of vacuolar invertase to stabilization of photosynthetic performance and energy metabolism was found to have a strong effect on whole plant physiology under stress.

## Results

### Stress-induced reprogramming of the primary metabolism

The multivariate analysis of stress-induced reprogramming of primary metabolism in both analysed natural accessions, C24 and Rsch, revealed that dynamics of absolute concentrations of citric acid, fumaric acid, sucrose, fructose and glucose most dominantly separated genotypes and/or conditions (Fig. 1; an overview of all loadings for PCA with and without autoscaling, i.e. zero mean-unit variance scale, is provided in Supplement 1a and 1b).

**Figure 1.**
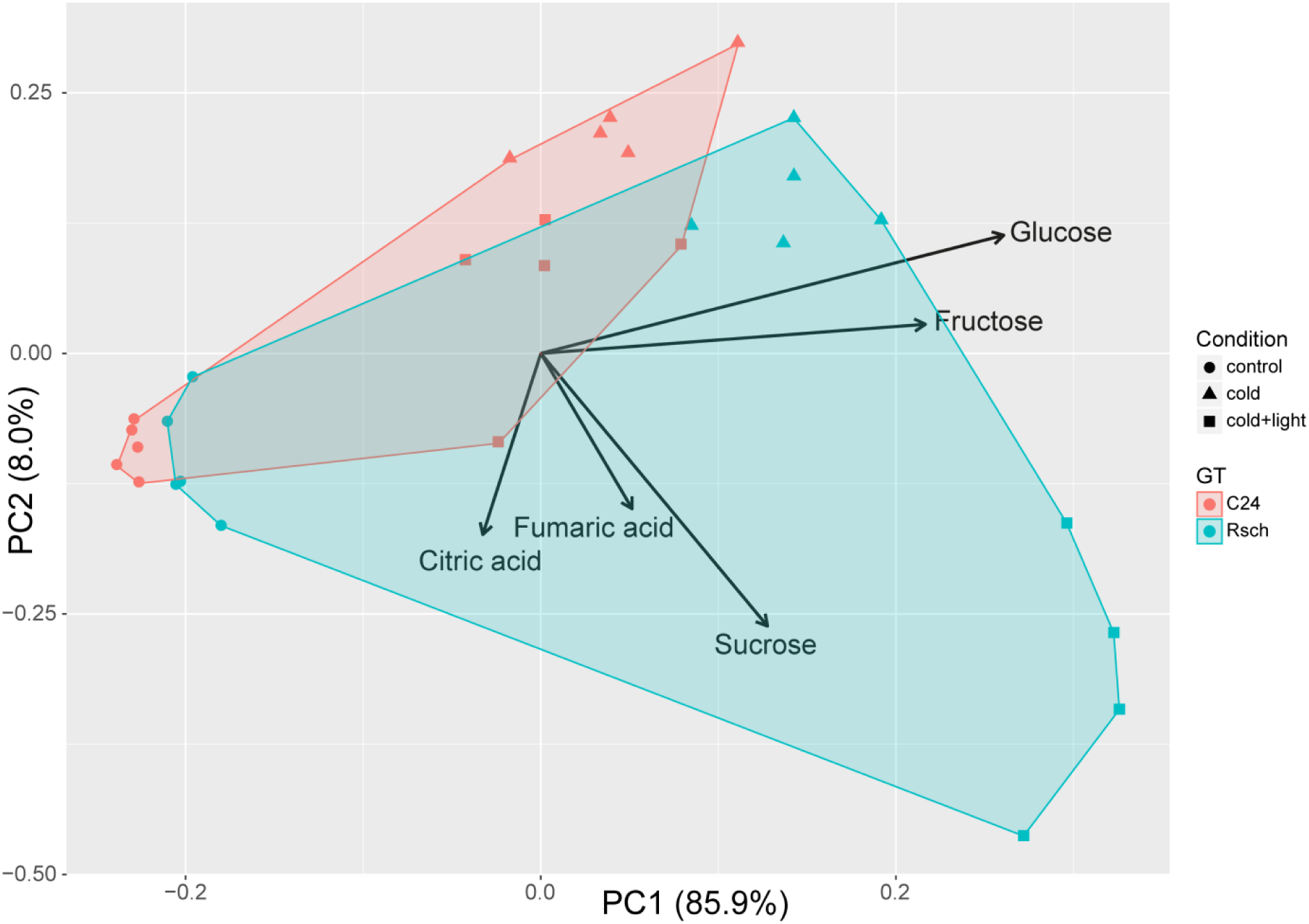
Principal component analysis of stress-induced reprogramming of primary metabolism. Samples of both natural *Arabidopsis* accessions are differently coloured. C24: red; Rsch: cyan. Different conditions are indicated by symbols. Filled circle: control; filled triangle: cold; filled square: cold and high-light. Arrows indicate the most dominant loadings, i.e. metabolites, separating accessions and conditions. PCA was performed using absolute metabolite concentrations without scaling.

While concentrations of citric acid were similar in both accessions under all considered conditions (Fig. 2 A), concentrations of fumaric acid significantly increased only in Rsch when exposed to combined stress conditions, i.e. cold and high-light (Fig. 2 B, ANOVA, p<0.01).

**Figure 2.**
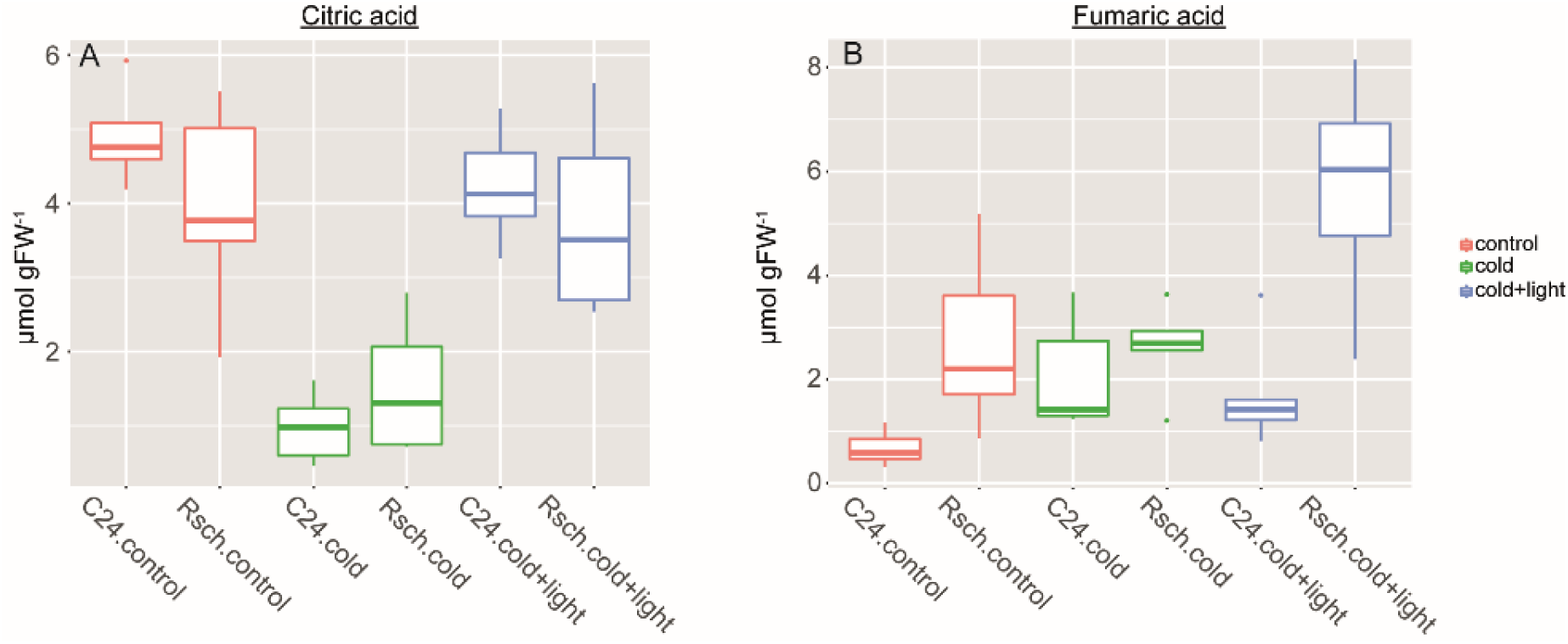
Stress-induced dynamics of the TCA cycle intermediates citric acid (A) and fumaric acid (B). Colours indicate conditions. Red: control; green: cold; blue: cold + high-light. Outliers are represented by dots (n≥4).

Similar to fumaric acid, also for sucrose, glucose and fructose the strongest increase was observed for Rsch plants under combined stress conditions (Fig. 3). While sugar concentrations also significantly increased in C24 in both stress conditions, this increase was always stronger in Rsch. When exposed to 5°C, sucrose concentration doubled in C24 while it increased three-fold in Rsch. When light intensity was increased from 80 to 320 μmol m^−2^s^−1^ within the cold, sucrose concentration in Rsch increased again three-fold (compared to control: 9-fold) while it stayed constant in C24 (compared to control: 2-fold) (Fig. 3 A). Also glucose and fructose concentrations showed a significant increase under low temperature always resulting in higher levels in Rsch than in C24 (Fig. 3B, C). While in Rsch both hexose concentrations increased significantly under combined stress conditions (p<0.01), in C24 no significant change could be observed and glucose concentration was rather decreasing (p=0.078) than increasing (Fig. 3 B). In summary, sucrose metabolism and fumaric acid showed a highly significant effect of genotype, condition and the combination of both factors (ANOVA, p<0.001) while citric acid was significantly affected only by low temperature (p<0.001).

**Figure 3.**
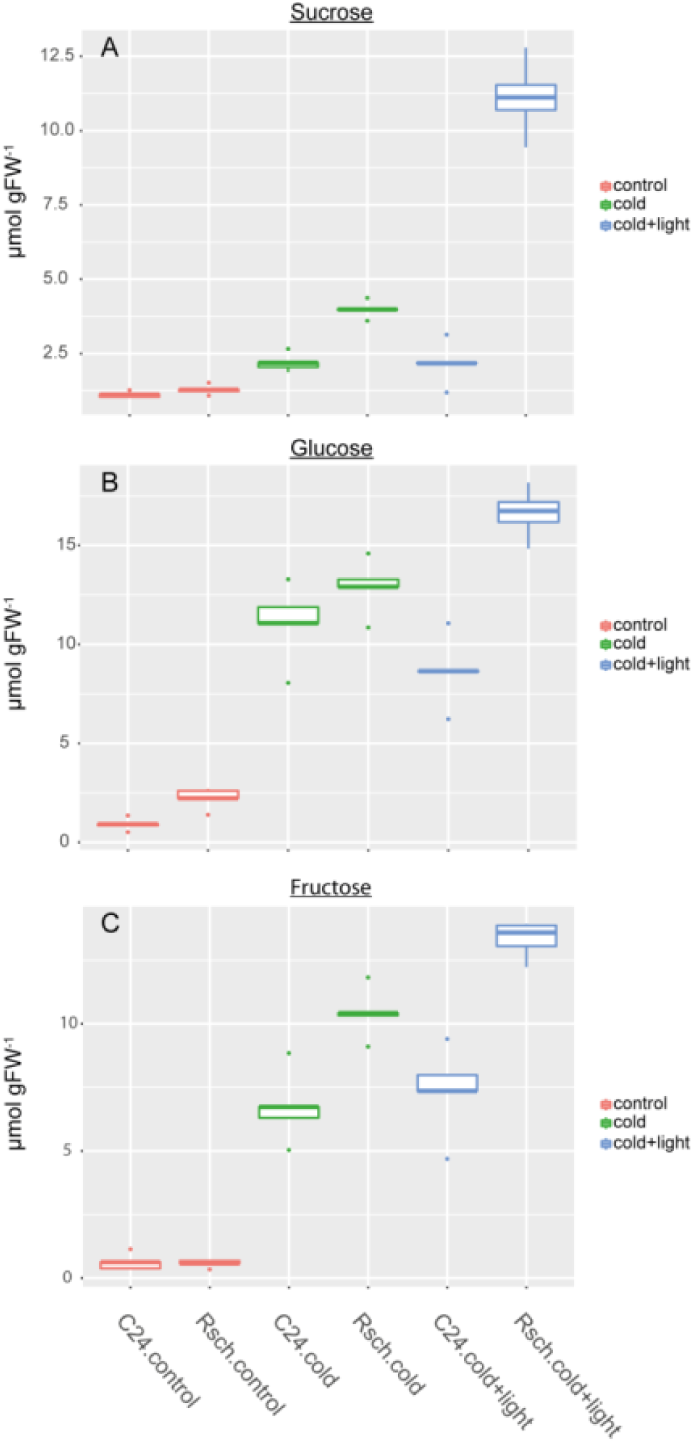
Stress-induced carbohydrate dynamics in C24 and Rsch. Boxplots are shown for sucrose (A), glucose (B) and fructose (C). Colours indicate conditions. Red: control; green: cold; blue: cold + high-light. Outliers are indicated by dots (n≥4).

### Subcellular distribution of sucrose cleavage

Due to the observed significant differences in sucrose and hexose dynamics between C24 and Rsch, the activity of neutral (cytosolic) and acidic (vacuolar) invertase was analysed because it is known to affect these sugar concentrations significantly (see e.g. (Klotke et al., 2004)). In addition, the subcellular distribution of invertase reaction substrate and product, i.e. sucrose, glucose and fructose, between plastids, cytosol and vacuole was determined applying a non-aqueous fractionation technique (Fürtauer et al., 2016).

The activity of neutral invertase showed a significant increase in C24 under combined stress condition while it only slightly increased after cold exposure (Fig. 4 A). In Rsch, no significant change of neutral invertase activity was observed across all conditions but activity was rather decreasing than increasing. In contrast, acid invertase activity of Rsch increased during cold exposure and reached a significantly higher level under combined cold and light stress (p<0.05; Fig. 4 B). In C24, acid invertase activity was significantly higher than in Rsch under all conditions (p<0.05). The activity peaked after 1 day of cold exposure and slightly decreased under combined stress conditions (Fig. 4 B).

**Figure 4.**
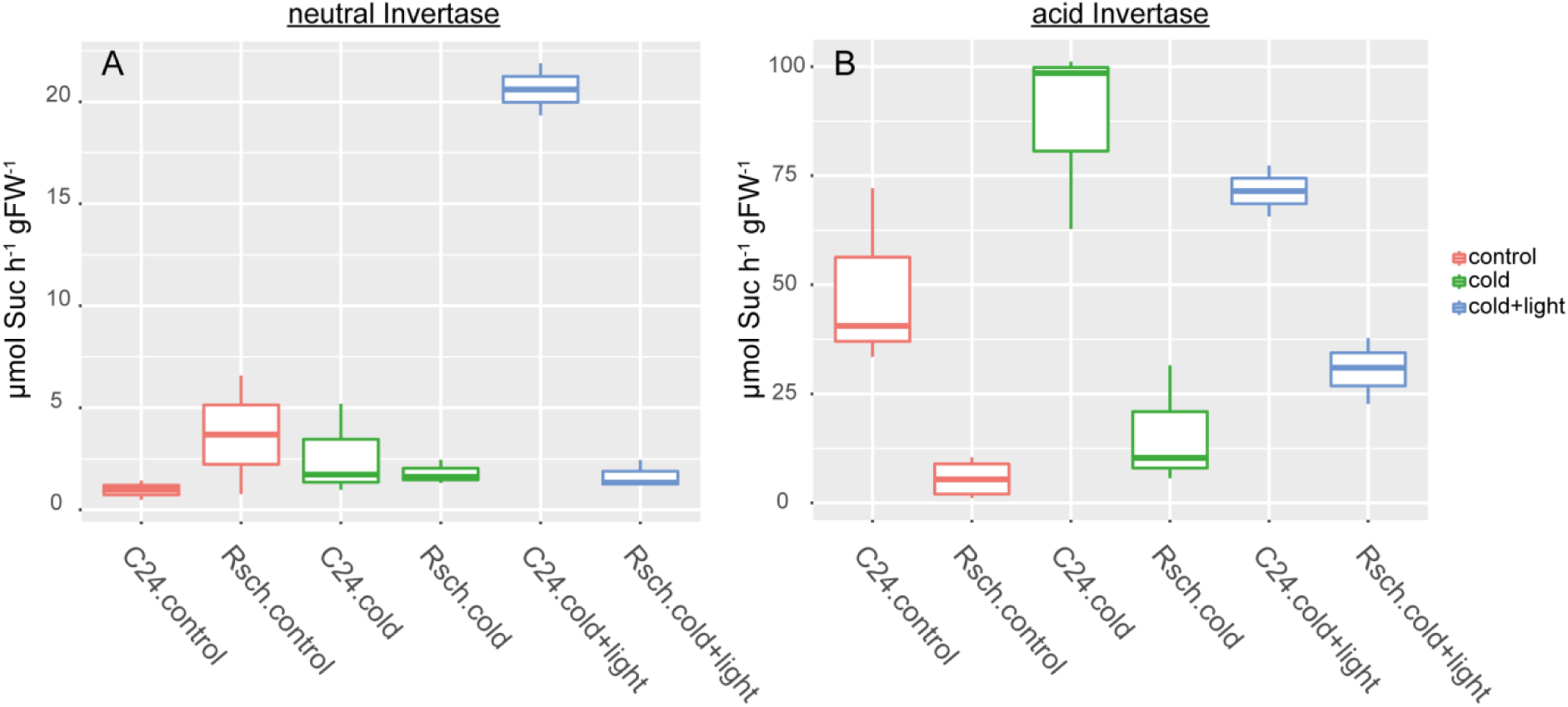
Invertase activity in C24 and Rsch under control and stress conditions. Neutral invertase activity (A) and acid invertase activity (B) were recorded under substrate saturation and were normalized to hour and fresh weight of the samples (n = 3). Colours represent conditions as indicated in the legend. Red: control; green: cold; blue: cold + high-light.

The subcellular distribution of sucrose, glucose and fructose showed strong stress-induced dynamics which differed between C24 and Rsch, particularly with respect to cytosolic and vacuolar distribution (Fig. 5). In both accessions, the plastidial concentration of all three sugars increased during cold and combined cold/light stress exposure and the increase was more pronounced in Rsch (Fig. 5 A, B). Similarly, cold exposure resulted in an increase of all sugars in the cytosolic and vacuolar compartment of both genotypes (Fig. 5 C-F). Yet, when cold exposure was combined with high-light exposure, cytosolic and vacuolar sugar concentrations in C24 either decreased (glucose) or stayed constant (fructose and sucrose; Fig. 5 C, E). In Rsch, combined stress resulted in a strong increase of vacuolar sugar concentrations as well as cytosolic sucrose concentration (Fig. 5 D, F). Hence, in summary, the combination of cold and high-light exposure uncoupled subcellular sugar reallocation in both accessions.

**Figure 5.**
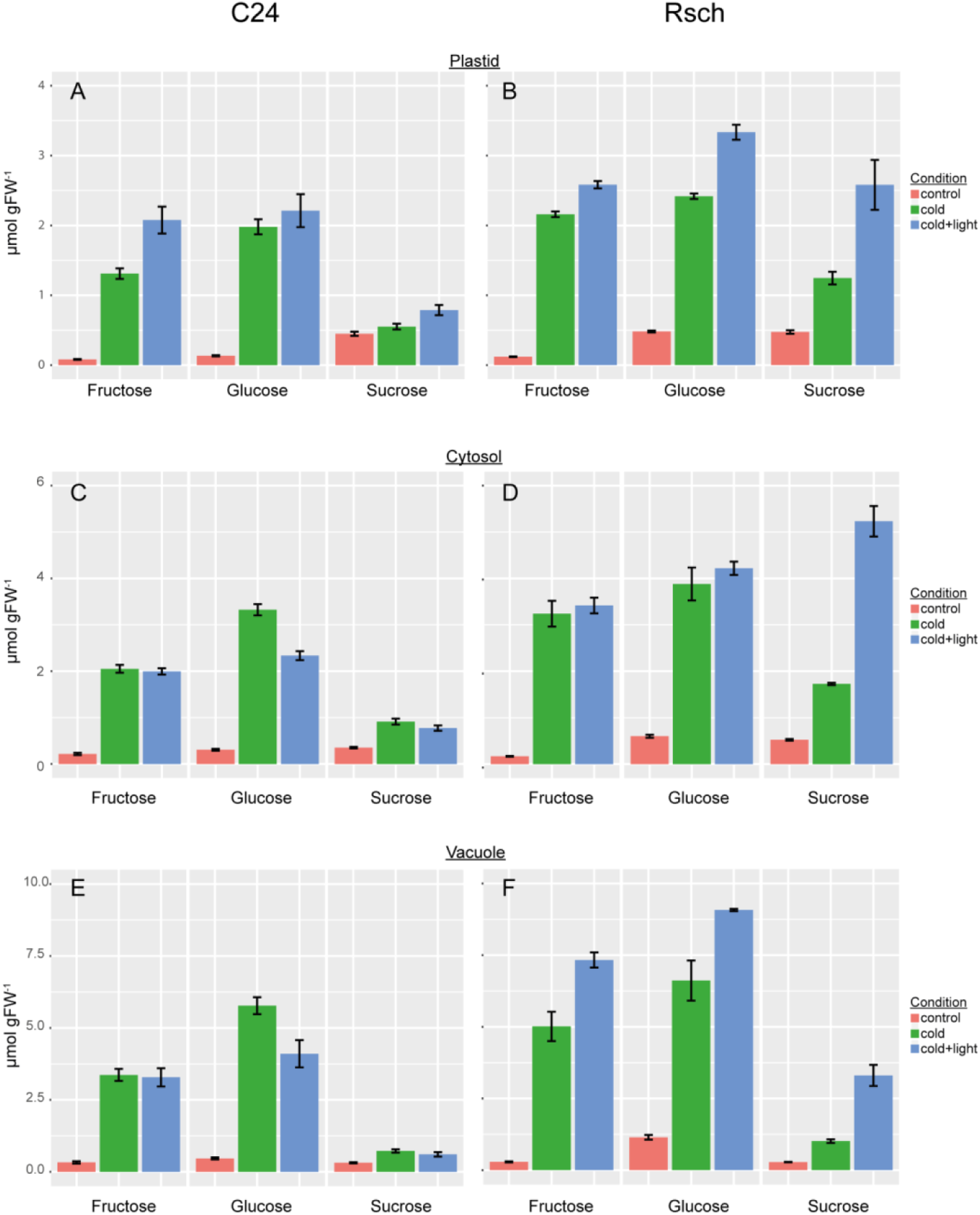
Subcellular sugar concentrations in C24 and Rsch during stress exposure. The left panel (A, C, E) represents concentrations in C24, the right panel (B, D, F) shows concentrations in Rsch. The upper panel (A, B) shows plastidial sugar concentrations, the middle panel (C, D) represents cytosolic concentrations and the lower panel (E, F) shows vacuolar concentrations. Colours represent conditions as indicated in the legend. Red: control; green: cold; blue: cold + high-light. Bars represent means ± SD (n = 4). Data on relative subcellular sugar distribution is provided in the supplements (Supplement 2).

### Kinetic modelling of stress-induced subcellular sugar allocation

To functionally combine experimental results of both enzyme kinetics and subcellular sugar concentrations, a mathematical model was developed to simulate reaction rates of sucrose cleavage within the cytosol (neutral invertase), the vacuole (acid invertase) and potential sugar transport between both compartments. The mathematical model comprised six ordinary differential equations describing dynamics in cytosolic and vacuolar concentrations of sucrose, glucose and fructose. For model simulation, experimentally determined subcellular sugar concentrations were used together with the kinetic parameters for invertase of this study as well of a previous study (Nägele and Heyer, 2013). To approximate *in vivo* kinetics of invertase enzymes in leaf metabolism at 22°C (control condition) and 5°C (stress conditions), measured enzyme activities were re-calculated using the Arrhenius equation as described earlier (Nägele et al., 2012). Detailed information about the model structure, kinetic parameters and Arrhenius equation is provided in the supplement (Supplements 3-5).

Simulation results revealed opposite trends of cytosolic and vacuolar sucrose cleavage in both considered accessions (Fig. 6). Already under control conditions, the mean ratio of vacuolar *versus* cytosolic invertase reaction rate was more than 20-fold higher in C24 resembling the experimentally determined ratio of acid and neutral invertase activity (see Fig. 4). Under cold and cold/light stress the mean ratio of simulated reaction rates clearly decreased in C24 (Fig. 6 A) while it increased in Rsch (Fig. 6 B).

**Figure 6.**
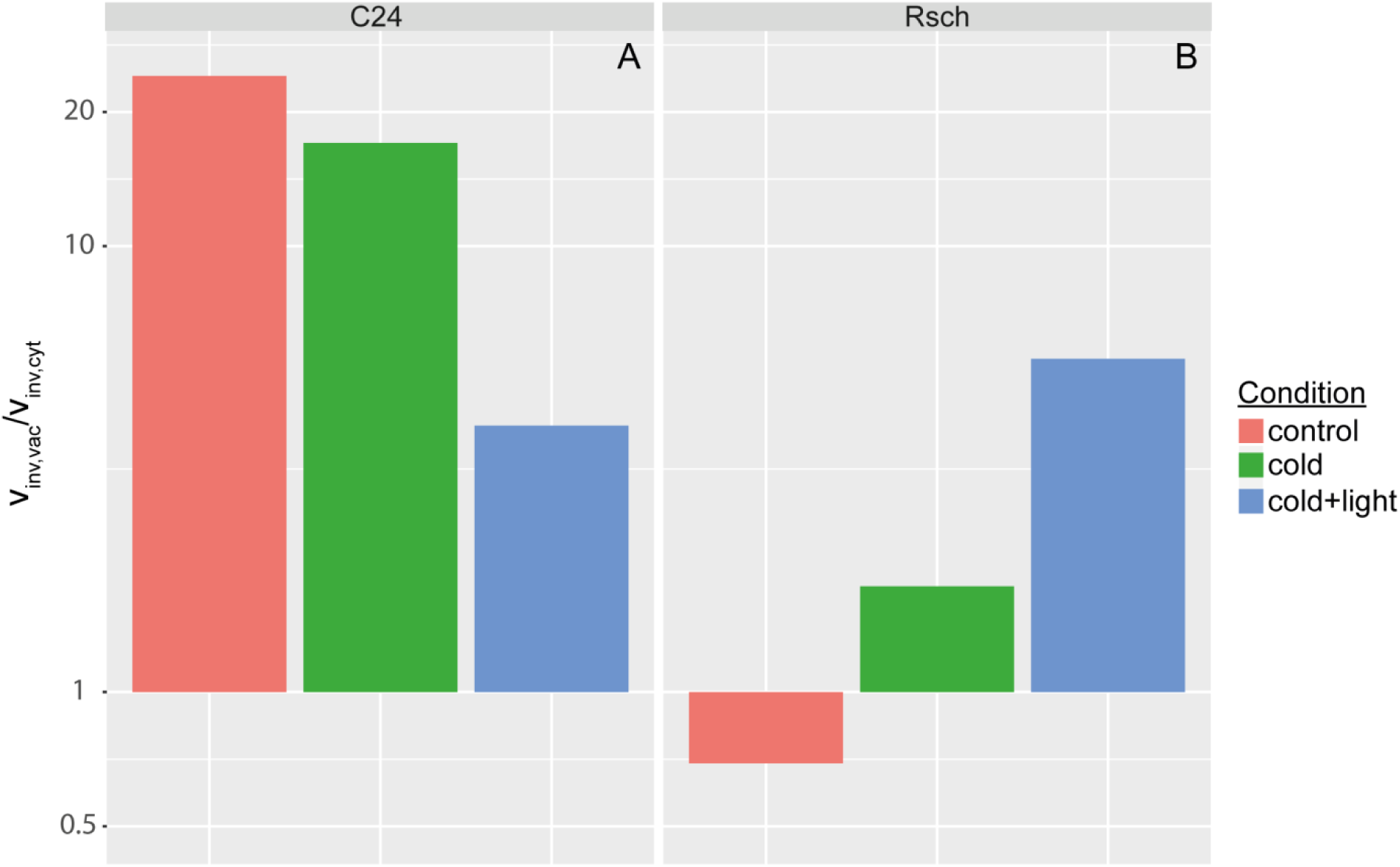
Ratio of vacuolar and cytosolic invertase reaction rates under all considered conditions. (A) Ratios of simulated reaction rates for C24, (B) ratios of reactions for Rsch. Ratios were built from mean values of simulations (n = 100).

Conclusively, not only the experimentally determined invertase activities, which were measured under optimum temperature and substrate saturation, showed a different stress-related pattern between C24 and Rsch, but also the simulation of thermodynamically adjusted subcellular reactions rates indicated a clearly different regulation of subcellular sugar metabolism during stress exposure.

### Invertase reactions contribute to photosystem stabilization during severe abiotic stress

Based on the observation that the more freezing-tolerant accession Rsch (Hannah et al., 2006) increases the proportion of vacuole sucrose cleavage during chilling stress (see Figure 4 and Figure 6), the role of vacuolar invertase activity during severe freezing and high-light stress was experimentally analysed. For this, plants of the T-DNA insertion line AtβFruct4 (SALK_100813; AT1G12240; *inv4*) were compared to wild type plants, Col-0. In *inv4*, vacuolar invertase activity was significantly reduced to ^~^3% of Col-0 (t-test p<0.01; Figure 7). Also neutral invertase activity was significantly reduced in *inv4* to ^~^30% of Col-0 (t-test p<0.05; Figure 7). Cell wall-associated invertase activity was not significantly affected in *inv4*.

**Figure 7.**
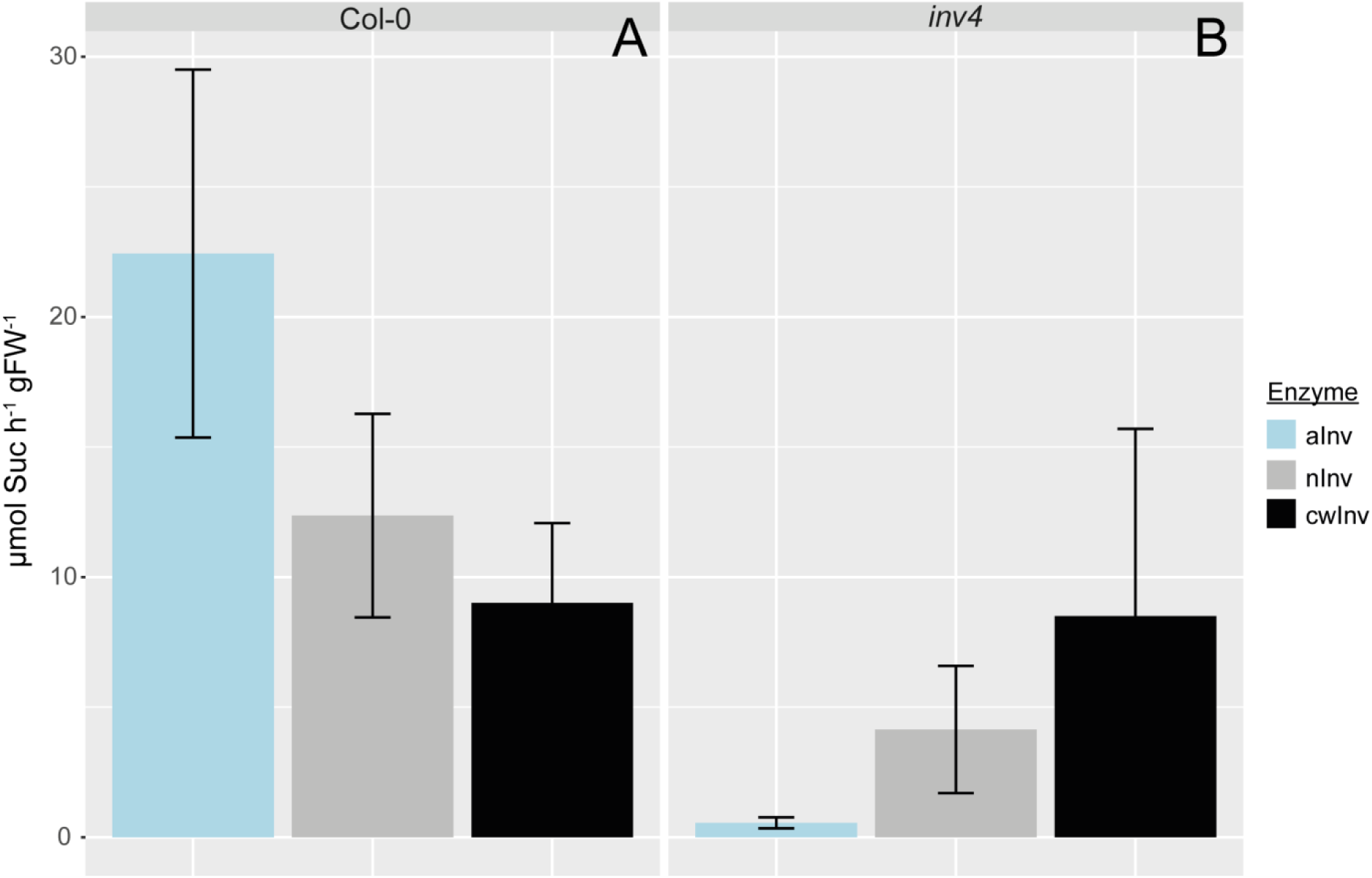
Activity of invertase isoforms in Col-0 and *inv4*. (A) Invertase activity of Col-0, (B) invertase activity of *inv4*. Colours indicate different isoforms. Cyan: acid invertase (aInv); grey: neutral invertase (nInv); black: cell wall-associated invertase (cwInv). Bars represent means ± SD (n=3).

From an ambient environment, i.e. a temperature of 20°C and light intensity of 100 μmol m^−2^ s^−1^, plants of both genotypes, i.e. Col-0 and *inv4*, and both natural accessions, C24 and Rsch, were transferred to freezing temperature (−20°C) and 500 μmol m^−2^ s^−1^ for 10 minutes. During this time period, the leaf temperature decreased to −2.5 ± 0.5°C (Fig. 8 C). This treatment was repeated three times and parameters of leaf chlorophyll fluorescence were recorded before as well as after 10, 20 and 30 minutes of stress exposure. Measurements were performed on dark adapted leaves at 20°C resulting in three freeze-thaw cycles.

**Figure 8.**
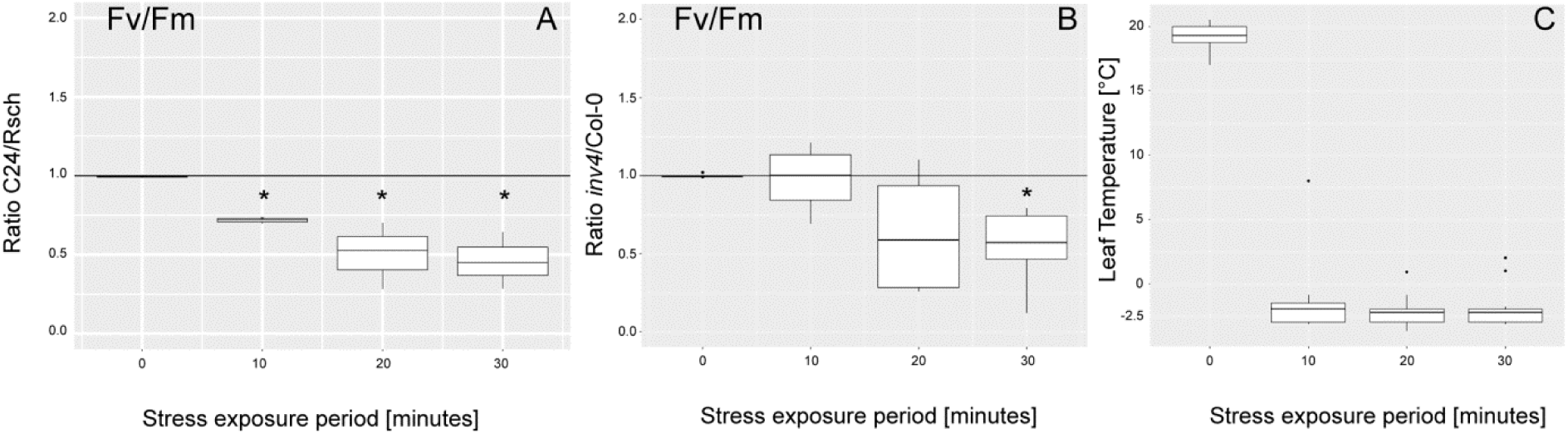
Ratio of the maximum quantum yield of photosystem II (F_v_/F_m_) during severe stress treatment. (A) Ratio of the F_v_/F_m_ parameter of C24 and Rsch, (B) Ratio of *inv4* and Col-0, (C) Leaf temperature during the combined freezing/high-light experiment (n = 5). Asterisks indicate significance (p<0.05). Boxplots represent 4 ratios which were measured pairwise.

During stress-application, the maximum quantum yield of photosystem II, F_v_/F_m_, decreased stronger in *inv4* than in Col-0 and stronger in C24 than in Rsch (Fig. 8 A, B). After 10 minutes, the F_v_/F_m_ - value of C24 was significantly lower than the value of Rsch (Fig. 8 A), and after 30 minutes it was significantly lower in *inv4* than in Col-0 (Fig. 8 B). This indicates that vacuolar and/or cytosolic invertase activity contributes to the stabilisation of photosynthetic performance under combined freezing and high-light stress. In addition, parameters of photochemical and non-photochemical quenching were determined using rapid light curves with an increasing photosynthetically active radiation (for details see *Materials and Methods*). While no genotype-specific difference was observed for C24 and Rsch, the non-photochemical quenching, which estimates the rate constant for heat loss from PSII, was significantly lower in *inv4* than in Col-0 after 30 minutes of stress application (ANOVA, p<0.001; see Supplement 6 and 7).

### Vacuolar invertase activity affects stress-induced reprogramming of the central energy metabolism

Potentially, stress-induced de-stabilization of the photosystem might be related to changes in energy metabolism due to effects on electron transport efficiency, proton motive force and ATP production. To prove the contribution of vacuolar invertase activity on cellular energy metabolism, levels of AMP, ADP and ATP as well as of sugar mono- and diphosphates were determined for Col-0 and *inv4* before and after 30 minutes of stress application. Results indicated a stress-induced increase of ATP concentration in both genotypes while this increase was ^~^3-fold stronger in *inv4* (Fig. 9 A). Further, AMP concentration in Col-0 decreased during stress while it remained stable in *inv4*. Concentration of ADP showed a slight stress-induced decrease in Col-0 while it significantly increased in *inv4*. Accompanying the stress-related increase of ATP concentrations, also the sum of glucose-6-phosphate and fructose-6-phosphate concentrations increased significantly in both genotypes (Fig. 9 B). The concentration of fructose-1,6-bisphosphate was not affected in Col-0 while it decreased in *inv4* (p=0.051) after 30 min of stress application.

**Figure 9.**
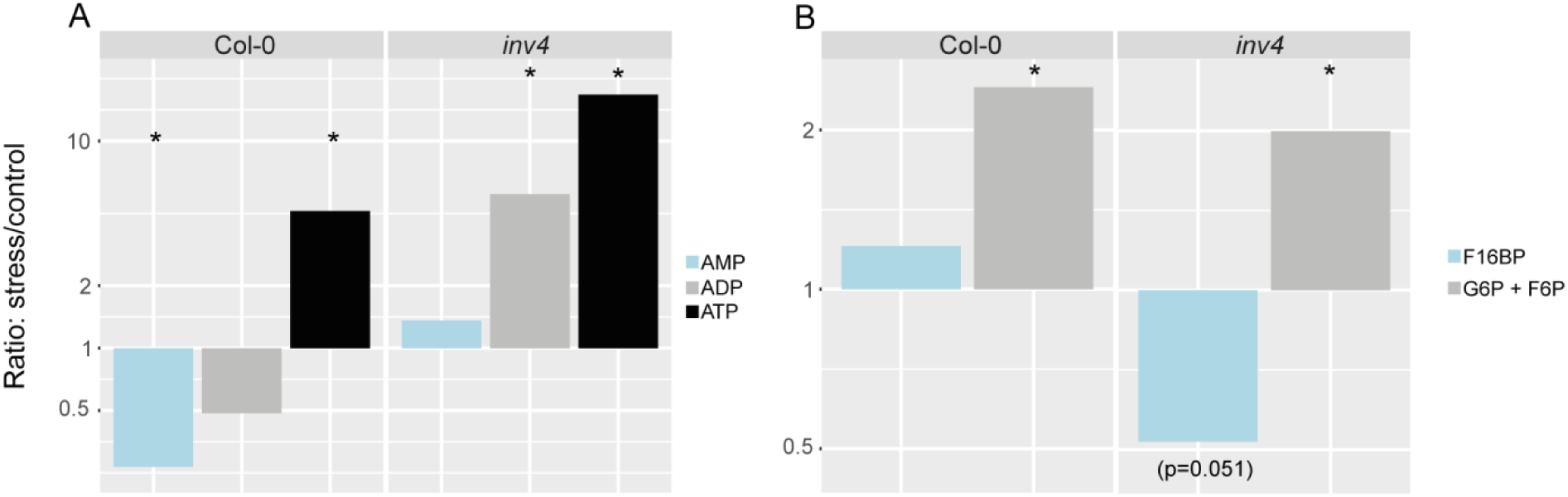
Stress effects in energy metabolism of Col-0 and *inv4*. (A) Stress-induced changes of nucleotide concentrations, (B) Stress-induced changes of sugar mono- and diphosphates. Ratios were built from mean values of concentrations determined for stressed (30 min; freezing + high-light) and control plants (n=3). Asterisks indicate significantly different mean values (p<0.05). AMP: Adenosine monophosphate; ADP: Adenosine diphosphate; ATP: Adenosine triphosphate; F16BP: Fructose-1,6-bisphosphate; G6P: Glucose-6-phosphate; F6P: Fructose-6-phosphate.

## Discussion

Regulation of carbohydrate metabolism is central to plant growth, development and stress response. As the direct output of photosynthetic CO_2_ fixation carbohydrates are substrate for biomass accumulation, biosynthesis of other organic molecules and play a central role in gene expression and signalling under abiotic stress (Gupta and Kaur, 2005; Sami et al., 2016). A characteristic response of many plants to cold stress is the fast and significant increase of soluble sugar concentrations which was also observed in the present study after 24h at 5°C in both analysed natural *Arabidopsis* accessions, C24 and Rsch. The increase of sucrose, glucose and fructose concentrations was stronger in the more cold and freezing tolerant Rsch accession which is in accordance with previous findings (Nägele and Heyer, 2013). Yet, under combined stress only Rsch continued to significantly increase sugar concentrations while concentrations in C24 were only slightly affected (sucrose and fructose) or even dropped again (glucose, see Fig. 3). Multivariate data analysis revealed a very strong impact on the separation of genotype and condition by sucrose dynamics (Fig. 1). A similar stress response could be observed for concentrations of fumaric acid where a strong concentration increase was only observed for Rsch under combined stress (Fig. 2 B). An explanation for these differences might be the recently described and naturally occurring promotor polymorphism of the *Arabidopsis* FUM2 gene (Riewe et al., 2016). The cytosolic fumarase, FUM2, catalyses the cytosolic interconversion of malate to fumarate and was shown to be essential for cold acclimation in *Arabidopsis* (Dyson et al., 2016). Riewe and colleagues found that the insertion/deletion polymorphism was absent from the promotor region of Columbia-0 *FUM2*, yet present in the C24 promotor region. Additionally, they provided evidence for the Col-0 *FUM2* allele in Rsch. Hence, as the insertion was found to be linked to reduced *FUM2* mRNA expression and fumarase activity (Riewe et al., 2016), this could explain the observed discrepancy of fumaric acid levels in C24 and Rsch and, as reported earlier, the differences in cold and freezing tolerance (Distelbarth et al., 2013). As citric acid concentrations showed completely different stress-induced dynamics (see Fig. 2 A) this indicates rather a specific effect for fumaric acid than a general stress response of TCA cycle intermediates.

### Subcellular partitioning of sucrose cleavage shapes photosynthetic performance under stress

Similar to fumaric acid, also subcellular sugar concentrations differed in dynamics and absolute values of both natural accessions, particularly under combined stress conditions (see Fig. 5). While in the cold tolerant accession Rsch all subcellular sugar concentrations increased both during cold and combined stress, the cytosolic and vacuolar sugar concentrations in C24 remained constant or even decreased under combined stress. This coincided with a dramatic shift of the vacuolar sucrose cleavage to the cytosol. The opposite was observed for Rsch where stress induced a shift of sucrose cleavage from the cytosol into the vacuole (Fig. 6). Together with the results from chlorophyll fluorescence measurements (Fig. 8) this indicates a stabilization of photosynthetic performance under stress by a shift of sucrose cleavage into the vacuole. Accordingly, compared to Col-0 photosynthetic performance under freezing and high-light stress was significantly reduced in the *inv4* mutant being strongly affected in the vacuolar sucrose cleavage capacity. While the accumulation of soluble sugars is well-known to be involved in cold stress response and freezing tolerance (see e.g. (Wanner and Junttila, 1999; Guy et al., 2008)), involved biochemical and regulatory mechanisms are less clear (Tarkowski and Van den Ende, 2015). Predominantly, this is due to diverse roles of sugars and involved enzymes in osmoregulation, cryoprotection, membrane stabilisation, signalling and ROS scavenging (Strand et al., 1999; Strand et al., 2003; Klotke et al., 2004; Heiber et al., 2014; Dahro et al., 2016). In previous studies, the sucrose biosynthesis pathway was shown to be a limiting factor in cold acclimation (Strand et al., 2003; Nägele et al., 2012). A balance between rates of carbon fixation and sucrose biosynthesis is a prerequisite for optimal rates of photosynthesis preventing the depletion of Calvin cycle intermediates or inorganic phosphate, P_i_, which would result in inhibition of ATP synthesis and Rubisco inactivation (Stitt and Hurry, 2002). Invertase catalyses the hydrolytic cleavage of sucrose and, thereby, strongly affects cellular sucrose concentration by degradation. As a consequence, invertase reaction, together with other sucrose-consuming processes like export to sink organs, substantially influences the physiologically relevant reaction equilibrium of sucrose biosynthesis in leaves by reducing the reaction product. Hence, potentially as a consequence of a deregulated sucrose biosynthesis/degradation equilibrium, the *inv4* mutant, which lacks
^~^97% of vacuolar invertase activity (see Fig. 7), was found to have reduced rate of net photosynthesis under ambient temperature (Nägele et al., 2010; Brauner et al., 2014).

If temperature drops below 0°C, intracellular ice formation might result in membrane lesions and extracellular ice nucleation causes cell dehydration (Steponkus, 1984). Several studies have provided evidence for a central role of invertase in response to drought and water stress (Kim et al., 2000; Andersen et al., 2002; Trouverie et al., 2003). In mature maize leaves, Trouverie and colleagues observed a fast and strong increase of vacuolar invertase and hexose levels after exposure to water deprivation (Trouverie et al., 2003). Another study identified a tonoplast-located monosaccharide transporter, ESL1, which was suggested to function in coordination with vacuolar invertase and to regulate osmotic pressure by sugar accumulation (Yamada et al., 2010). While those findings point to a role of invertase activity in regulating cellular osmotic pressure during drought, it is unclear if it has a similar function in cold and freezing stress. Previously it was reported that inhibition of vacuolar invertase had no effect on basic freezing tolerance of non-acclimated plants of the accessions C24 and Col-0 (Klotke et al., 2004). Further, this study showed that increased sucrose concentrations and sucrose-to-hexose ratios are not sufficient to improve basic freezing tolerance. While those findings are related to freezing tolerance information gained from electrolyte leakage measurements and, hence, do not reflect the experimental setup in the present study it still rather questions vacuolar invertase to adjust osmotic pressure during acute low temperature stress. This is further supported by the finding that during early low temperature response the cold susceptible accession C24 has a higher vacuolar invertase activity (Fig. 4 B) than the tolerant accession Rsch, yet being significantly less freezing tolerant (Nägele et al., 2011). Conclusively, a separate interpretation of stress-induced dynamics of sucrose and hexose concentrations and invertase activity hardly provides a conclusive and mechanistic picture of subcellular stress response of carbohydrate metabolism. In contrast, kinetically connecting substrate and product concentrations with enzyme activity to simulate subcellular *in vivo* fluxes reveals a more realistic picture of sucrose-hexose ratios in context of metabolism. The ratio of simulated vacuolar and cytosolic invertase fluxes revealed a strong decrease during cold and combined stress conditions in the susceptible accession while in the tolerant accession the ratio clearly increased (see Fig. 6). The increasing proportion of vacuolar sucrose cleavage in Rsch may provide an explanation for its stronger increase of cytosolic sucrose concentration when compared to C24: while in Rsch cytosolic invertase activity was found to decrease during stress, it increased in C24, particularly under combined stress condition (see Fig. 4 A). Cytosolic sucrose is substrate for other stress-relevant metabolites, e.g. for raffinose (Schneider and Keller, 2009). Further, cytosolic sucrose is involved in signalling processes, ROS scavenging and membrane stabilization being central to abiotic stress response and acclimation (Tarkowski and Van den Ende, 2015). In summary, a stress-induced shift of sucrose cleavage from the cytosol into the vacuole might enable cytosolic sucrose accumulation being essential for a coordinate and efficient plant stress response.

### Vacuolar invertase deficiency affects the plastid-cytosol interaction during stress response

Stabilization of energy metabolism is a central element of cellular stress response (Baena-Gonzalez and Sheen, 2008). Thioredoxin together with enzymes being involved in its metabolism, for instance NADPH thioredoxin reductase C (NTRC), have been identified to be core components of a regulatory network coordinating involved pathways, e.g. photosynthesis, TCA cycle and mitochondrial respiration (Gelhaye et al., 2004; Meng et al., 2010; Daloso et al., 2015; Thormählen et al., 2015). Recently, different environmental stress conditions were observed to affect the redox state of only a particular set of organelles resulting in the idea of organelle redox autonomy (Bratt et al., 2016). Similar to the experiment in the present study, Bratt and co-workers analysed seedlings of *Arabidopsis thaliana* exposed to freezing temperature, high-light and a combination of both. Under combined stress, they observed the strongest change in oxidation degree in chloroplasts which were affected ^~^5-fold stronger than mitochondria whereas almost no effect could be quantified for cytosol and peroxisome (Bratt et al., 2016). This allows us to speculate that the observed stress-induced changes of nucleotide levels in the present study were (predominantly) related to redox dynamics in the chloroplast (Fig. 9 A). While for Col-0 a decrease of AMP and ADP was observed, ATP concentration rose about 5-fold due to stress application indicating a substrate depletion of photosynthetic ATP generation. While ATP levels of *inv4* mutants were also found to increase, yet about 3-fold stronger than in Col-0, also ADP levels increased and AMP levels were slightly elevated by stress exposure. Together with the finding of reduced F16BP concentrations, lower F_v_/F_m_ value than in Col-0 and a similar stress response of hexose phosphate concentrations, this indicates a deregulated chloroplast-cytosol interaction in *inv4* during stress. In illuminated leaves of *Arabidopsis*, 70% of F16BP were previously found to be located in the chloroplast and 30% in the cytosol (Szecowka et al., 2013). Within the same study, more than 60% of G6P and F6P were found to be located in the cytosol indicating a dominant stress-effect of vacuolar invertase deficiency rather in the chloroplast than in the cytosol.

A possible scenario explaining these findings is the reduction of invertase-driven hexose generation under freezing temperature which is more dramatic in *inv4* knockout mutants than in the wild type. Previously, we have shown that the hexokinase-driven rate of hexose phosphorylation is reduced in *inv4* (Nägele et al., 2010), most probably resulting in a reduced (cytosolic) ATP turnover and futile cycling of sucrose. This affects the translocation of triose phosphates into the cytosol, the transport of P_i_ into the chloroplast and inhibits the usage of energy absorbed by PSII for ATP synthesis (Stitt and Hurry, 2002). As a consequence, stronger imbalance of photosynthetic primary and secondary reactions in *inv4* results in earlier ROS generation during stress exposure causing photoinhibition and a faster decrease of F_v_/F_m_ than in the wild type (see Figure 10).

**Figure 10.**
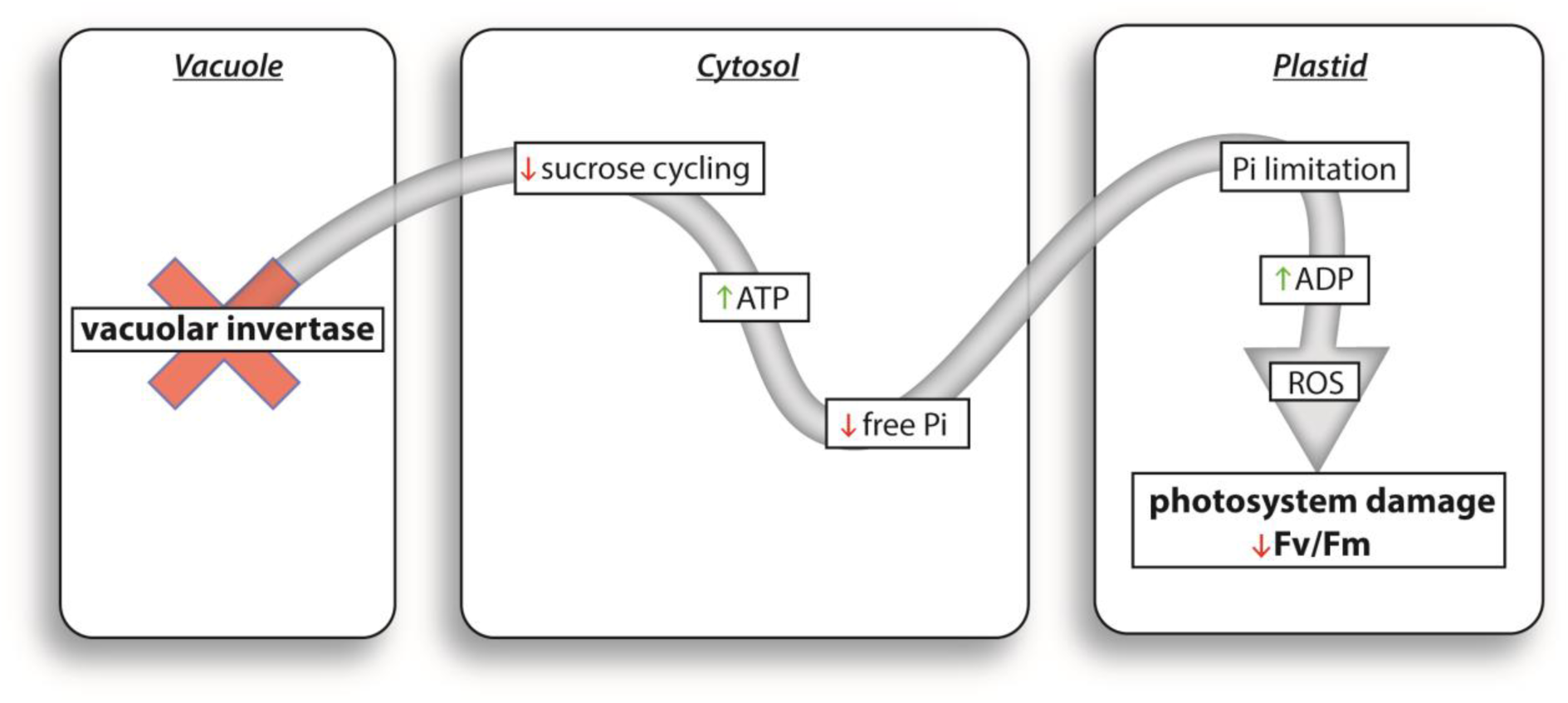
Schematic representation of the suggested effect of vacuolar invertase activity on photosynthesis and energy metabolism. ADP: Adenosine diphosphate; ATP: Adenosine triphosphate; P_i_: inorganic phosphate.

## Material and Methods

### Plant Material

Plants of *Arabidopsis thaliana* accessions Columbia (Col-0), C24 and Rschew (Rsch) as well as of the T-DNA insertion line *At*β*Fruct4* (SALK_100813; AT1G12240; *inv4*) were grown within a growth cabinet under fully controlled conditions (Conviron^®^ Adaptis). Light intensity under control condition was 80 μmol m^−2^ s^−1^ in a 12/12h day/night cycle. Relative air humidity was 60% and temperature was 22°C during the day and 18°C during the night. All plants were grown on soil which was composed of Einheitserde^®^ ED63 and perlite. Plants were watered daily and fertilized once with NPK fertilization solution (WUXAL ^®^Super; MANNA°-Dünger, Ammerbuch). For the cold stress experiment, plants of Rsch and C24 were exposed to 5°C for 24 hours. For the combined cold-light stress experiment, plants were subsequently exposed to a 4-fold higher light intensity, i.e. 320 μmol m^−2^ s^−1^, at 5°C for 3 hours. All samples were collected at the middle of the light phase, i.e. 6 hours after light on, except for the samples of the combined stress experiment which were taken after 9 hours in the light phase (6hours cold + 3 hours cold/light). Each sample consisted of three leaves from 3 different plants which were grown in three different pots. Samples were immediately quenched in liquid nitrogen and stored at −80°C until experimental analysis.

For the combined freezing and high-light experiment, plants of Col-0 and *inv4* were exposed to −20°C and a light intensity of 500 μmol m^−2^ s^−1^. The light source was a LED-panel. The actual leaf temperature (-2.5 ± 0.5°C, Fig. 8 C) was estimated using an infrared thermometer (Basetech Mini 1).

### Chlorophyll fluorescence measurements

Parameters of chlorophyll fluorescence were recorded using a WALZ MINI-PAM II/B (Heinz Walz GmbH, Germany). For measurement, the genotypes Col-0 and *inv4* as well as C24 and Rsch were subjected to the severe stress condition (-20°C and 500 μmol m^−2^ s^−1^) in parallel. The first measurement, i.e. control measurement, was performed with plants at room temperature after dark adaptation over 5 minutes. Then, every 10 minutes during the stress application, a plant was dark adapted for 3 minutes at −20°C and a “Rapid Light Curve” was recorded at 22°C over 5 minutes. The intensities of photosynthetically active radiation (PAR) were 0, 24, 43, 63, 87, 122, 185, 280, 411, 619, 808, 1130, and 1773 μmol m^−2^ s^−1^ which were each applied for 20 seconds.

### Non-aqueous fractionation of leaf material

Subcellular metabolite levels were determined applying the non-aqueous fractionation technique as described previously (Fürtauer et al., 2016). Finely ground and lyophilized leaf material was resuspended in a mixture of tetrachlorethylene (C_2_Cl_4_) and n-heptane (C_7_H_16_). As described in the protocol (Fürtauer et al., 2016), a density gradient of this mixture was applied to separate the specific compartment densities within the suspension. To determine the distribution of the compartments across the gradient, compartment-specific marker enzyme activities were determined and correlated with the relative metabolite levels. Marker enzymes were pyrophosphatase (plastid), UDP-glucose pyrophosphorylase (cytosol) and acid phosphatase (vacuole). This revealed the relative subcellular metabolite distribution across the compartments. To determine the absolute compartment-specific metabolite concentration, values of relative distribution were multiplied with the absolute total metabolite concentration determined by GC-ToF-MS analysis (see next paragraph).

### GC-ToF-MS Analysis

Primary metabolite concentrations were absolutely quantified via gas chromatography coupled to time-of-flight mass spectrometry applying a previously published protocol with slight modifications (Weckwerth et al., 2004; Nukarinen et al., 2016). Frozen leaf samples were ground to a fine powder using a ball mill (Retsch GmbH, Haan, Germany). Primary metabolites were extracted twice with a methanol-chloroform-water mixture (MCW, 5/2/1, v/v/v) followed by an extraction step with 80% ethanol in which the samples were heated to 80°C for 30 minutes. Water was added to the MCW supernatant to induce a phase separation. The polar phase was then mixed with the ethanol extract and dried in a vacuum concentrator (ScanVac, LaboGene). The dried extracts were derivatized applying methoximation (Methoxyamine hydrochloride in pyridine) and silylation (N-Methyl-N-(trimethylsilyl) trifluoroacetamide). For methoximation, samples were incubated for 90 minutes at 30°C. For silylation, samples were incubated for 30 minutes at 37°C. Derivatized samples were transferred into glass vials which were sealed with a crimp cap. GC-ToF-MS analysis was performed on an Agilent 6890 gas chromatograph (Agilent Technologies^®^, Santa Clara, USA) coupled to a LECO Pegasus^®^ GCxGC-TOF mass spectrometer (LECO Corporation, St. Joseph, USA). Compounds were separated on an Agilent column HP5MS (length: 30 m, diameter: 0.25 mm, film: 0.25 μm). Deconvolution of the total ion chromatogram and peak integration was performed using the software LECO Chromatof^®^. For absolute quantification, calibration curves were recorded for each metabolite comprising 5 different concentrations within the linear range of detection.

### IC-MS Analysis

Nucleotides and phosphorylated sugars were extracted using a cooled 80% methanol solution. Samples were extracted twice and supernatants were pooled and filtered through a 10μm polyethylene filter (MobiSpin Column “F”, MoBiTec GmbH, Goettingen, Germany) prior to measurement. Concentrations of nucleotides and phosphorylated sugars were determined on a Dionex^™^ ICS-4000 Capillary HPIC system (Thermo Fisher Scientific, Waltham, Massachusetts, USA) coupled to a triple-quadrupole mass spectrometer (TSQ Vantage, Thermo Fisher Scientific, Waltham, Massachusetts, USA). Substances were chromatographically separated within a KOH gradient (2mM to 120mM) with a flow rate of 0.015 ml min^−1^ and a total gradient duration of 25 min (Column: Dionex IonPac^™^ AS11-HC-4μm, RFIC^™^ Capillary, 0.4×250mm). Relative concentration was determined by integrating peak areas of selected reaction monitoring (SRM). Mass transitions and collision energies for SRM were chosen according to previously published protocols (see e.g. (Heinzel and Rolletschek, 2011)). Retention time and SRM settings of AMP, ADP, ATP, G6P and F6P were validated using chemical standards (Sigma-Aldrich, St. Louis, USA). The F16BP peak was not validated by a chemical standard. Peak areas were quantified using the Thermo Xcalibur software (Version 2.3.26; Thermo Fisher Scientific, Waltham, Massachusetts, USA).

### Measurement of enzyme activities

Activity of neutral and acid invertase was measured as described previously with small modifications (Nägele et al., 2010). In brief, about 100 mg of frozen and finely ground powder of leaf material was suspended in 50 mM HEPES-KOH (pH 7.4), 5mM MgCl_2_, 1mM EDTA, 1mM EGTA, 1mM phenylmethylsulfonylfluoride (PMSF), 5mM dithiothreitol (DTT), 0.1% Triton-X-100 and 1% glycerine. Suspensions were centrifuged with 21,000g at 4°C for 5 minutes and the supernatant was transferred to a new reaction tube. Both invertase activities were determined in this supernatant. Soluble acid invertase activity was determined in 20 mM Na-acetate buffer (pH 4.7) and 100 mM sucrose as substrate. Neutral invertase was assayed in a 20mM HEPES-KOH (pH 7.5) buffer and 100 mM sucrose as substrate. The control, i.e. blank, sample was heated to 95°C for 3 minutes before the addition of plant extracts. All reactions were incubated for 20 minutes at 30°C and stopped by boiling for 3 minutes. The glucose concentration of each sample was determined photometrically.

### Mathematical modelling and statistics

Mathematical modelling and numerical simulation of sucrose metabolism was performed within the software environment MATLAB^®^ (www.mathworks.com) using the SBPOP package (http://www.sbtoolbox2.org/main.php) (Schmidt and Jirstrand, 2006). The mathematical model consisted of six ordinary differential equations (ODEs) describing time-dependent changes in concentrations of sucrose (Suc), glucose (Glc) and fructose (Frc) both in the cytosolic (cyt) as well as the vacuolar (vac) compartment. A graphical representation of the model structure is provided in the supplements (Supplement 3). All model reactions were assumed to follow Michaelis-Menten kinetics (Eq. 1), except for the input flux (modelled by constant flux) as well as the output flux (modelled by mass-action kinetics).

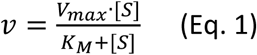

Here, *ν* denotes the reaction rate, *ν_max_* is the maximum enzyme activity determined under substrate saturation, *K_M_* represents the Michaelis-Menten constant, i.e. enzymatic substrate affinity, and [*S*] is the substrate concentration. Reaction products of sucrose cleavage, i.e. the free hexoses glucose and fructose, inhibited invertase activity competitively (fructose) and non-competitively (glucose) (Sturm, 1999). The experimentally determined *ν_max_* values were adjusted to the treatment temperature (control: 22°C; stress: 5°C) applying the Arrhenius equation as described earlier (Nägele et al., 2012). Further, all kinetic parameters were estimated using a combination of global and local parameter optimization. In a first optimization step, kinetic parameters were estimated using a particle swarm pattern search algorithm for bound constrained global optimization (Vaz and Vicente, 2007). In a second step, the model containing the globally optimized parameter set was subject to a fit analysis, i.e. a repeated perturbation and optimization to yield information about reproducibility depending on the parameter values. This second optimization step was performed in 100 replicates using a downhill simplex method in multi-dimensions (Press, 2002). Results of this fit analysis are provided in the supplement (Supplement 4).

Statistical analysis was performed with R (https://www.r-project.org/) and R Studio (https://www.rstudio.com/).

## Acknowledgments

We would like to thank the members of the Mosys Department for critical discussion and helpful comments on the manuscript. Particularly, we thank the gardeners Andreas Schröfl and Thomas Joch for their support and expertise in plant cultivation. We also thank Gert Bachmann and Wolfgang Postl for supporting our study with assistance in PAM-measurements and their self-constructed LED-panel. Further, we thank Matthias Nagler for excellent advice and helpful comments within the presented topic. This work was supported by the Austrian Science Fund (FWF; Project I 2071), and the Vienna Metabolomics Center ViMe at the University of Vienna.

## Author contributions

JW performed kinetic modelling. JW, LF and TN performed experiments, evaluated the data and wrote the paper. WW contributed materials and methods. TN conceived the study.

## Supplements

### Supplement 1a: PCA data not scaled

Interactive PCA plot including all variables, i.e. primary metabolites, without scaling (if problems occur with opening the file, please try a different web browser or change your browser settings).

### Supplement 1b: PCA data scaled

Interactive PCA plot including all variables, i.e. primary metabolites, after autoscaling. Metabolite concentrations were scaled by subtracting the mean value and dividing by the standard deviation, i.e. zero mean-unit variance (if problems occur with opening the file, please try a different web browser or change your browser settings).

### Supplement2:NAF relative distribution

Relative subcellular distributions of glucose (Glc), fructose (Frc) and sucrose (Suc) in plastid (Pla), cytosol (Cyt) and vacuole (Vac) in the three examined conditions, determined by non-aqueous fractionation and GC-ToF-MS (see Methods).

### Supplement3: Model graph

Graphical representation of the ODE model used for kinetic modeling. The model comprises enzymatic (blue) and transport (black) reactions as well as product inhibition of both invertase isoforms (red arrows). Suc: sucrose, Glc: glucose, Frc: fructose, inv: invertase, cyt: cytosolic, vac: vacuolar

### Supplement4: Parameter boxplots

Fitted and scaled parameters from 100 perturbation / optimisation runs. (A) C24 control, (B) C24 cold stress, (C) C24 cold + high-light stress, (D) Rsch control, (E) Rsch cold stress, (F) Rsch cold + high-light stress. G: Glucose, F: Fructose, inv: invertase, cyt: cytosol, vac: vacuole, Km: Michaelis-Menten constant, Ki: inhibition constant, k: activity constant, vmax: maximum enzyme activity, t: transport, out: efflux reation, in: influx reaction

### Supplement5: Arrhenius corrections

Maximum enzyme activities were measured at 30°C. To account for the dependency of the reaction rate on the temperature prevalent in the experimental setup, an adjustment of measured reaction rates through the Arrhenius’ Equation (Eq. 2) was conducted.

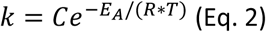

Where C is the frequency factor of the reaction and E_A_ denotes the activation energy. As the exact activation energy is unknown, the mean of several calculations using typical E_A_ values ranging from 40 to 50 kJ/mol was taken (Bisswanger, 2008).

### Supplement6: NPQ

Non photochemical quenching (NPQ) levels over the course of the RLC measurement at 0, 10, 20 and 30 minutes of combined cold and high-light stress. The lines are loess-type smoothing splines with a 95% confidence interval indicated by the grey area.

### Supplement7: qP

Photochemical fluorescence quenching coefficent (qP) levels over the course of the RLC measurement at 0, 10, 20 and 30 minutes of combined cold and high-light stress. The lines are loess-type smoothing splines with a 95% confidence interval indicated by the grey area.

